# When offspring outsmarting parents: Neuronal genes expression in two generations of marine parasitic worm

**DOI:** 10.1101/2023.06.17.545403

**Authors:** Oleg Tolstenkov, Marios Chatzigeorgiou, Alexander Gorbushin

## Abstract

Trematodes, or flukes, cause disease in millions of people, impact animal health, and alter the functional organization of biological communities. During the transition from the intramolluscan redia to the free-living cercaria stage in a complex life cycle of trematodes, extensive anatomical and behavioral modifications occur, enabling the cercaria to locate and infect the next host in the complex water environment. However, the functional changes that occur in the nervous system during this shift are not well understood.

We used a *de novo* transcriptome to characterize the molecular building blocks of the trematode nervous system and identify pathways that may underlie differences in nervous system function between the rediae and cercariae stages of the *Cryptocotyle lingua*, marine trematode species causing problems for fisheries. Our results confirmed the streamlined molecular toolkit of these parasitic trematodes, including the absence of certain key signaling pathways and ion channels. We documented the loss of nitric oxide synthase not only in *C. lingua* but also in the entire phylum Platyhelminthes. We identified several neuronal genes upregulated in dispersal larvae, including genes involved in synaptic vesicle trafficking, TRPA channels, G-protein coupled receptors, and surprisingly nitric oxide receptors soluble guanylate cyclase. Validation of these findings using neuronal markers and *in situ* hybridization allowed us to hypothesize the protein function in relation to the adaptations and host-finding strategy of the dispersal larva. Our results and established behavior quantification toolkit for cercaria motility provide a foundation for future research on the behavior and physiology of parasitic flatworms, with potential implications for developing antiparasitic measures.

**Highlights:**

- We utilized a behavior quantification toolkit and described essential neuronal genes in a handy model species, enabling the study of fluke neurobiology at the systems level.
- We characterized and validated neuronal gene expression profiles in cercarial embryos within rediae and swimming host-searching cercariae.
- The streamlined molecular toolkit of parasites reveals the absence of important signaling pathways and ion channels in their nervous system.
- We documented loss of nitric oxide synthase in flatworms.
- The expression pattern of nitric oxide receptors, soluble guanylate cyclases, upregulated in swimming larvae, emphasizes their crucial involvement in the dispersal process.
- Two upregulated TRPA channels in cercaria are primarily expressed in cilia and peripheral neurons, emphasizing their importance in host finding.

## Introduction

Trematodes, or flukes, cause disease in millions of people, impact animal health, and alter the functional organization of biological communities. To develop strategies to combat these severely detrimental effects, it is fundamental to understand their biology, especially that of the nervous system, through research on different model species across life stages ^1^. Enabled by earlier work on the genomic and transcriptomic analyses of a few model trematode species of medical relevance, some major features of neuronal gene organization in trematodes have already been characterized ^2, 3^. However, limiting research to only a few trematode species is disadvantageous because trematodes are known to exhibit species-dependent host-pathogen interactions, which in turn give rise to highly variable mechanisms of host finding and infection. An urgent challenge in the field is to broaden the spectrum of trematode species that researchers work on. Thus, this work aims at contributing to the expansion of marine trematode models for neuroscience research, such as to those that are not hazardous to humans and thus allowing for more quantitative and systematic experimentation in the laboratory setting.

Nervous system streamlining and plasticity are fundamental processes that shape neural function over time, enabling the nervous system to adapt to changing environmental demands. Trematodes have a highly complex life cycle that few animal models can match. For instance, the generational transition from the intramolluscan redia to the freely swimming cercaria comes with extensive anatomical and behavioral modifications, enabling the larva to locate and infect the next host in the complex water environment ^4, 5^. However, the functional changes that occur in the nervous system during this transition are not well understood.

The trematode species *Cryptocotyle lingua* (Creplin, 1825) (Digenea: Heterophyidae) is widespread in the North Atlantic Ocean, in European waters and the Northwest Atlantic (WoRMS database). It uses coastal birds as definitive hosts, where the parasites develop as hermaphroditic adults and produce eggs. A ciliated larval stage, called miracidia, hatches from the egg and infects sea snails (periwinkles, genus *Littorina*), where it develops into a sporocyst. The latter produces parthenogenetic daughter generations represented by rediae ^6^. The rediae produce and release free-swimming, short-lived and eye-spotted cercariae that demonstrate characteristic sensitivity to light and water turbulence stimuli ^7, 8^. These larvae infect downstream hosts (Figure 1 A), causing the common „black spot disease‟ in several marine fish species including the cod, which can cause loss to both the economy and the ecosystem ^9^. Here, cercariae penetrate the fish skin, encyst in the hypodermis, and survive as metacercariae for a long time until the intermediate host is eaten by the definitive host. Aside from causing significant problems for fisheries ^10–13^, *C. lingua* is often used in ecological ^14, 15^, immunobiological ^16^ and behavior studies ^17, 18^.

**Figure 1.**
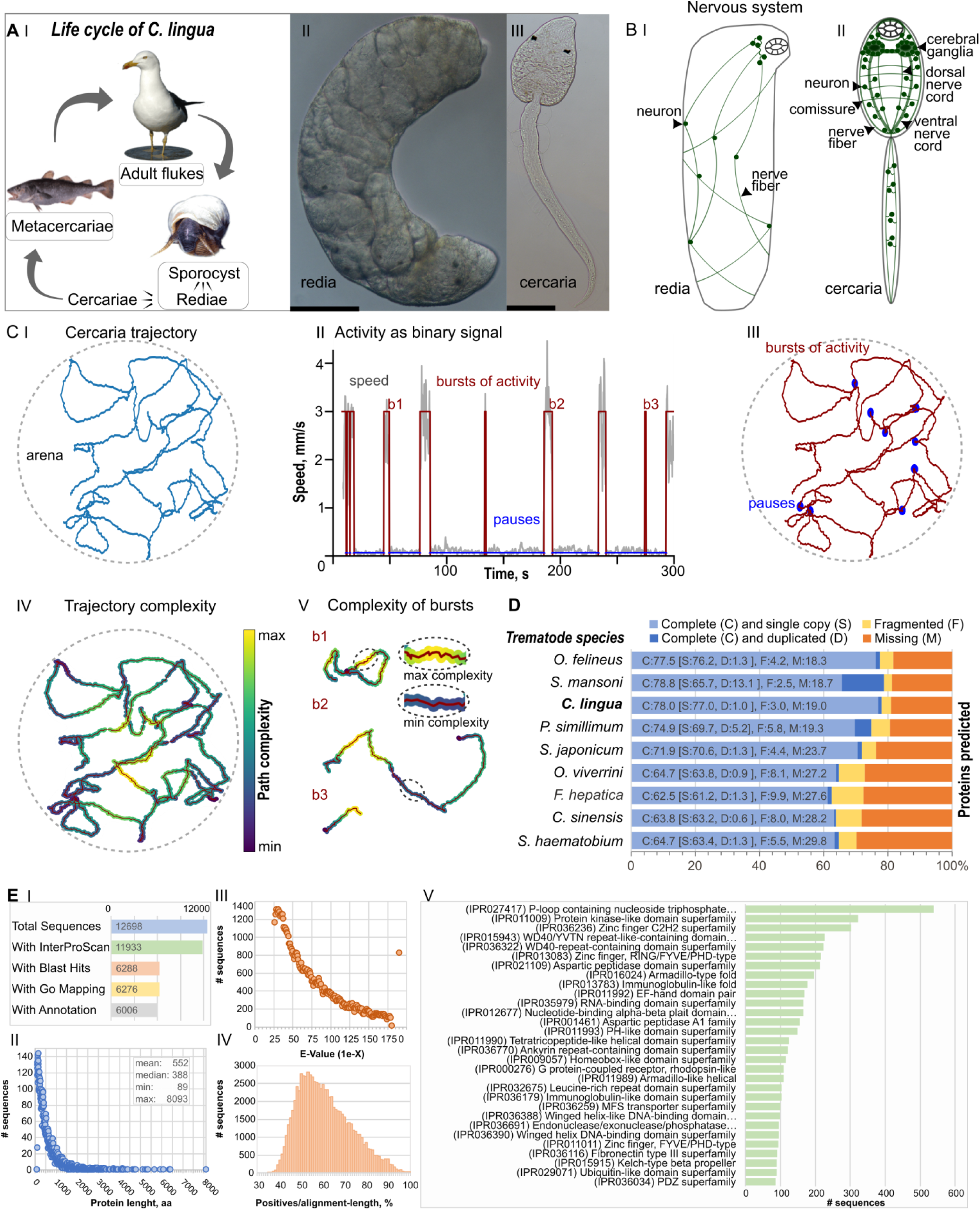
Introduction in the model species *C. lingua*, transcriptome assembly and protein prediction. **(A)** *C. lingua* life cycle (I), intramolluscan redia with developing cercariae embryos (II) and free-swimming cercaria (II), scale bar = 50 µm. **(B)** Nervous system increases anatomic complexity in cercaria (II) compared to redia (I), cartoon based on the neuronal markers staining (see below). **(C**) Trajectories analysis showed complexity of cercaria behavior. Representative trajectory (I), traditional approach of cercaria behavior analysis reveals easy to quantify two states of activity (II) that are plotted on trajectory (III), and results of path complexity (entropy) analysis with complexity of trajectory with 1 second window (IV) and individual bursts of activity with specified parts of maximum and minimum path complexity (V). **(D)** Summarized BUSCO (v4.1.4; Metazoa_odb10) benchmarking of protein arrays predicted in *C. lingua* transcriptome and in proteomes of eight other digenean species (*Schistosoma mansoni* (UP000008854), *S. japonicum* (UP000311919), *S. haematobium* (UP000471633), *Opisthorchis viverrine* (UP000054324), *O. felineus* (UP000308267), *Clonorchis sinensis* (UP000286415), *Fasciola hepatica* (UP000230066) and *Psilotrema simillimum* ^24^. Bar charts show proportions of 954 BUSCO orthologs classified as complete, fragmented, and missing. **(E)** Descriptive statistics of proteins inferred from *C. lingua* transcriptome. (I) summary of protein statistics; (II) length distribution of predicted proteins; (III) E-value distribution of BLASTp hits in SwissProt database; (IV) protein sequence similarity to curated SwissProt prototypes; (V) top 30 InterProScan protein domain superfamilies.

Despite the importance of the nervous system in the trematode life cycle progression, little is known about the molecular mechanisms underlying this process. To address this gap in knowledge, we conducted a transcriptomic analysis of the nervous system in cercariae and rediae of the fluke *C lingua*, focusing on genes differentially expressed between these two phases of the life cycle. To elucidate the molecular and behavioral basis of the promising model species *C. lingua* for further neurobiological studies, this work aims to gain insights into nervous system function in two stages of the life cycle. To establish *C. lingua* as an experimental model in functional neuroscience, this work has characterized the complexity of behavior of the cercaria stage using live behavioral recording combined with computer vision. Furthermore, to link these animal behaviors to neural function, nervous system development and gene expression, we performed *de novo* transcriptome assembly of *C. lingua* across two life-cycle stages, in combination with whole-mount antibody staining and *in situ* hybridization analysis of gene expression patterns, to identify important genes in neurotransmission that may play cercariae stage-specific functions.

## Results

### Dispersal larvae of *C. lingua* exhibit complex but quantifiable behavior

The complex life cycle of trematodes requires successful transmission between multiple hosts and depends on the ability of the dispersal larva (cercaria) to quickly locate a suitable host within the host-space environment ^19–21^. In contrast, rediae are tissue parasites that proliferate and produce cercariae (Figure 1A) in the host’s circulatory system ^6^, gonads and in hemolymph lacunae ^22^. The redia has a rudimentary nervous system ^23^, that is specifically adapted to the intramolluscan environment. In contrast, fully developed cercariae have a functional nervous system that allows them to sense and respond to environmental cues ^5^ (Figure 1B). In our experiments, we could not qualify spontaneous slow wiggling of rediae in ASW as behavior (Supplemental movie (Sm1). Cercariae of *C. lingua*, consistent with previous studies, exhibited intermittent swimming behavior (Supplemental movie Sm2, and Figure 1C). However, based on our analysis of path complexity during spontaneous locomotion, our data reveals previously uncharacterized patterns within burst-like activity in *C. lingua* cercariae. This suggests that their behavior may exhibit a higher degree of complexity and potentially operate in more dimensions than previously thought (Figure 1 C I-V).

Deep sequencing of *C. lingua* transcriptome resulted in high quality assembly and protein prediction. We obtained 159.1 million of strand-specific paired reads in all six libraries (21.1 – 23.9 million per each) which were used in the ’raw’ transcriptome assembly. The fraction of the raw assembly, the ’curated assembly’, was produced by filtering out non-coding Trinity clusters (’genes’), low-expressed genes and few genes of xenogenic origin (the host *L. littorea*, bacteria, fungi and human). Out of 12698 proteins, 6288 were annotated with a BLAST hit (E-value cut-off: 1E-25) and sequence descriptor. Predicted protein length varied from 89 to 8093 amino acids. BUSCO completeness assessments and protein statistics showed high assembly quality and protein prediction comparable to other trematode model species ^24^ (Figure 2 D, E).

**Figure 2.**
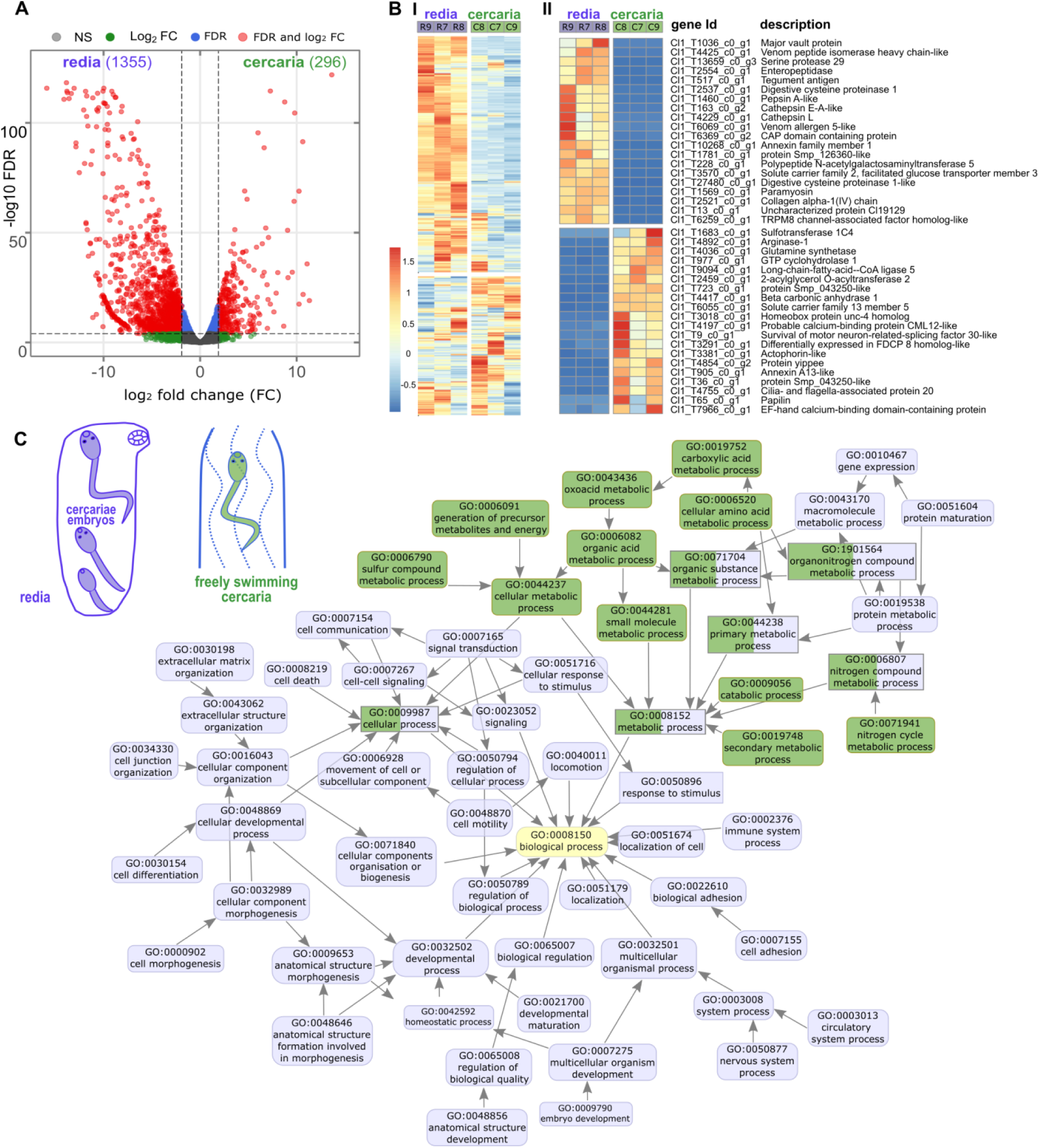
Differential expression analysis of *C. lingua* transcriptome (9338 unigenes) was consistent with the production and dispersal functions of redia and cercaria. (A) Volcano plot of differentially expressed genes (DEGs) in rediae and cercariae. Numbers of stage specific DEGs are shown in parentheses. The red dots indicate significantly regulated genes (*threshold*: logFC >= 2, FDR < 0.05), the blue and green dots indicate under threshold genes, respectively for FC and for FDR, the grey dots indicate non-significant difference. **(B)** Heatmap showing gene expression profiles (cross-sample normalized TMM values) in three redia and three cercaria samples. Red and blue indicate up- and down-regulated expression (I). The scale bar represents the row Z-score. (II) Top 20 proteins that were significantly down- and up-regulated in cercariae. **(C)** Combined graph of Gene ontology (GO) term enrichment analysis for differentially expressed genes in *C. lingua* rediae and cercariae. Only overrepresented (FDR < 0.05) functional annotations for Biological processes are displayed. Annotations are indicated with nodes colored by developmental stage of the parasite.

### Gene expression patterns reveal redia’s role in cercaria production and highlight cercariae’s dispersal-related characteristics

The transcriptome of *C. lingua* contains a high number of mobile elements, which we excluded from our analyses. In the subset of 9338 protein-coding genes, 1355 (14.5%) were four-fold upregulated (FDR≤0.05) in rediae, while only 296 (3.2%) in cercariae (Fig. 2 A, B). Among top 20 stage-specific genes, several proteinases required for enzymatic digestion were presented in redia-specific DEGs, consistent with the active feeding of intramolluscan redia. The main biological function of redia is to produce dispersal larvae cercariae, and not surprisingly, only one of 74 development regulatory Homeobox genes, protein unc-4 homolog, involved in motor neuron determination and maintenance ^25^, was upregulated in cercaria. Meanwhile, 28 Homeobox genes were specifically expressed in redia stage highlighting a putative homebox signature underlying *C. lingua* embryonic development (Fig 2B).

To gain insight into the stage-specific biological processes in the *C. lingua* expression profile, we conducted a Gene Ontology (GO) enrichment analysis on the DEG sets of rediae and cercariae. The results were consistent with the expected biology of these life cycle stages: the parthenogenetically breeding generation (rediae) produces actively swimming larvae (cercariae). Rediae, which contain all stages of cercaria embryo development, expressed genes that control various cellular processes, including cell communication, developmental processes, cell differentiation, and anatomical structures, such as morphogenesis and developmental maturation (Fig. 2C). These features are characteristic of the parthenogenetic generation (redia) (Fig. 2C). Additionally, redia-specific gene expression included multicellular organismal processes that regulate digestion and nervous system functions. In contrast, mature cercariae were sampled after actively swimming for at least four hours after shedding and GO analysis revealed a highly represented group of terms related to metabolic processes, including catabolic processes, primary, cellular, and small molecular metabolism (Fig. 2C). This finding is consistent with the dispersal function of endotrophic larvae, which use internal sources of energy for swimming and indicates the high metabolic rate in host seeking larva cercaria. Notably, the rediae exhibited a different metabolic process, which was represented by protein and macromolecular metabolism (Fig. 2C).

### Neural signatures in transcriptomes of rediae and cercariae highlighted production of significant number of upregulated proteins at redia stage

To investigate the molecular-level function of the nervous system in *C. lingua*, we focused on 364 referenced neural proteins, identified and classified 229 homologous protein-encoding transcripts in the transcriptome of *C. lingua* (NCBI GenBank, accession numbers MW361065 – MW361239). We performed a differential expression analysis of selected neuronal genes in redia and cercaria and characterized the domain architecture of the protein and the expression profile of each neuronal gene (Supplement information file and Supplement figures 1 and 2). We characterized transcripts involved in chemical and electric synaptic machinery, transmission with classical neurotransmitters and small gaseous messengers, reception machinery, voltage gated channels, and genes encoding sensory proteins (Supplement information file and Supplement figures 1 and 2, Supplement tables 1-8). Here we shortly characterize the sets of neuronal proteins and their expression between stages.

#### Neurotransmitters biosynthesis and metabolism

The biosynthesis of classical neurotransmitters (NT) and small gaseous messengers is a crucial step in neurotransmission, and the identification of corresponding enzymes in the transcriptome provides evidence for the existence of these pathways ^26^. We confirmed presence of biogenic amines pathways: dopamine (tyrosine 3-monooxygenase (TH), aromatic-L-amino-acid decarboxylase (DDC)), norepinephrine (dopamine beta-hydroxylase), tyramine (tyrosine decarboxylase (TDC), serotonin (tryptophan 5-hydroxylase 1); acetylcholine (choline O-acetyltransferase, acetylcholinesterase), glutamate (glutaminase) and transsulfuration pathway (cystathionine beta-synthase) The complete list of enzymes and more details can be found in (Fig. 3 A; Suppl. Table 1, Supplement information file and Supplement Figure 1 A).

**Figure 3.**
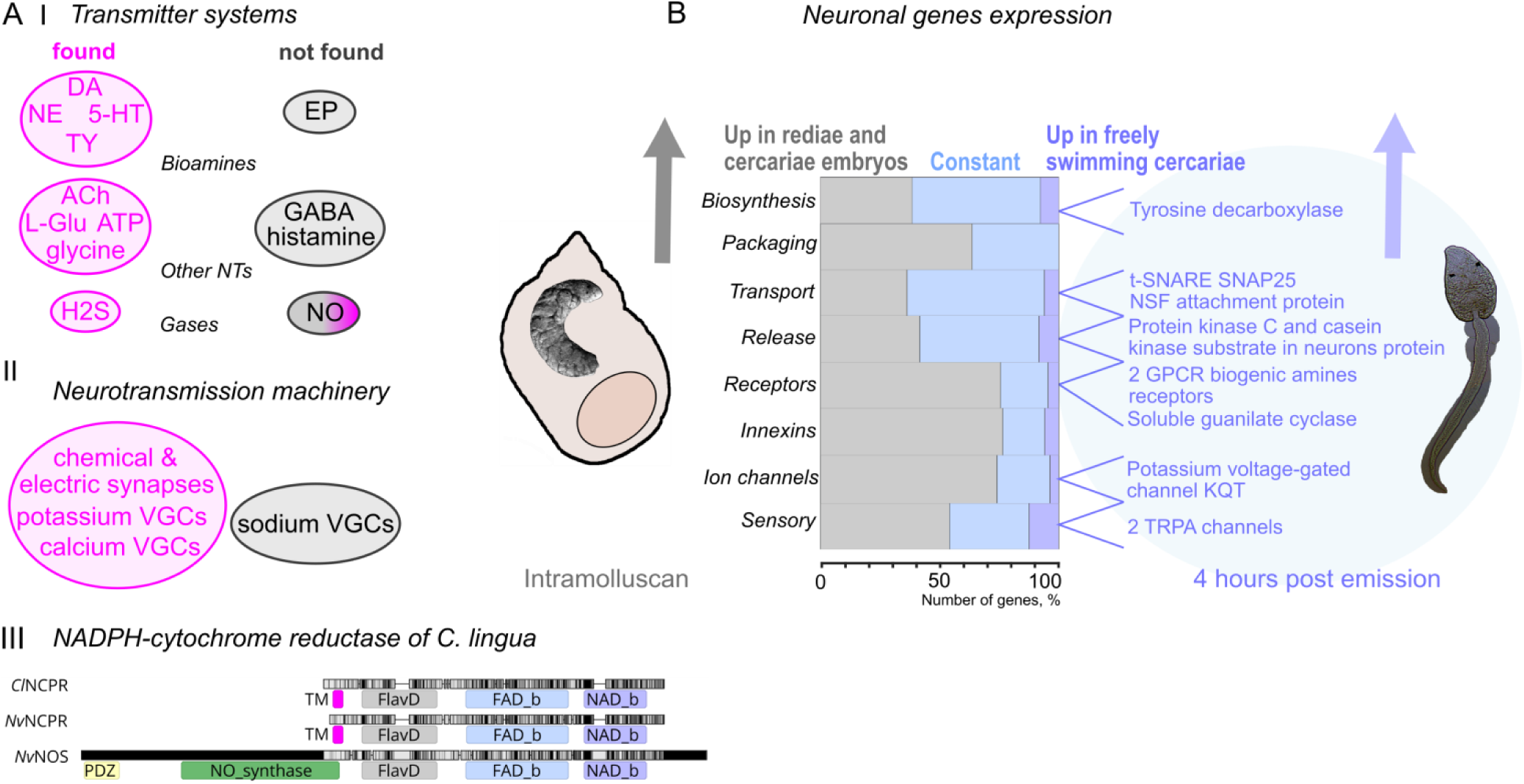
Neuronal proteins are not equally expressed in rediae and cercariae. (**A**) Presence (magenta) and absence (grey) of homologs proteins related to neurotransmitter systems (**I**) and machinery of neurotransmission (**II**). (I) Based on enzymes involved in biosynthesis, specific proteins of neurotransmitter packaging, receptors and methabolism we registered the neurotransmitters pathways for dopamine (DA), norepinephrine (NE), serotonin (5-TH), tyramine (TY), acetylcholine (Ach), glutamate (L-Glu), glycine, hydrogen sulfide (H2S), and adenosine triphosphate (ATP). We did not register homologues of proteins involved in epinephrine biosynthesis (EP), pathways of gamma-aminobutyric acid (GABA), histamine, and nitric oxide syntases (NOS). While the nitric oxide (NO) receptors soluble guanylate cyclases were present. (II) Presence and absence of varios voltage gated channels (VGCs). (**III**) Domain alignment of *C. lingua* NADPH--cytochrome P450 reductase (*Cl*NCPR), closest protein to NOS in *C. lingua* transcriptome with two *Nematostella vectensis* enzymes from CPR family, the orthologous protein *Nv*NCPR (XP_032236207) and Nitric oxide synthase, brain isoform X1 *Nv*NOS (XP_032236179). PFAM domains PDZ, NO_synthase, Flavodoxin, FAD-, NAD-binding and transmembrane helices (TM) are shown. (**B**) Representation of neuronal DEGs in two stages studied. Genes upregulated in rediae are shown in grey, constant in blue and the genes upregulated in freely swimming larvae 4 hours after shedding are in lilac blue. Percentage was calculated from the number of genes in each subgroup characterized. Differential expression (p<0.05) was calculated for three paired biological replicates (See methods section).

Only a few genes from this group were significantly upregulated in rediae, including orthologs of biogenic amine synthesis, such as TH and DDC, as well as enzymes that are not specific to the nervous system, such as cholinesterases and glutaminase. TDC was the only gene upregulated in cercaria. The expression levels of the majority of the genes did not change significantly in rediae and cercariae (Figure 3B).

We failed to find orthologs of several genes involved in neurotransmitter biosynthesis and metabolism, including phenylethanolamine N-methyltransferase, tyramine beta-hydroxylase, DBH-like monooxygenase protein 1, cytochrome P4502D6, cmine oxidase [flavin-containing] A, catechol O-methyltransferase, amine oxidase [flavin-containing] B, histidine decarboxylase, histamine N-methyltransferase, glutamate decarboxylase 1 and 2, heme oxygenase 1, and nitric oxide synthase 1, 2, and 3 in *C. lingua* transcriptome.

The nitric oxide synthase (NOS) enzyme belongs to a family of CPR (cytochrome P450 Reductase) enzymes that contain both flavodoxin-like (FMN-binding) and FAD domains. This family includes NADPH-cytochrome P450 reductase (NCPR) and bacterial sulfite reductase ^27^. In the *C. lingua* transcriptome, only NCPR (GeneBank: MW361191) is present, which is composed of four structural motifs: transmembrane helices, flavodoxin-like domain, FAD-binding domain, and NAD-binding domain (Fig. 3 C). However, there are no transcripts in the assembly encoding protein with the NO_synthase (syn. NOS_oxigenase, NOSoxy) domain, which is the hallmark structural characteristic of all types of NOS ^28^. Moreover, this N-terminal and functionally central domain is absent in all 33 Platyhelminthes genomes and 49 transcriptomes (NCBI TSA projects) available, including representatives of classes Trematoda, Monogenea, Cestoda, and free-living turbellarians *Macrostomum lignano* and *Schmidtea mediterranea* (BLASTp and tBLASTn searches in WormBase Parasite v. WBPS14).

#### Synaptic vesicles loading

Consistent with the NT systems our analysis revealed the presence of genes involved in vesicle loading, including vesicular acetylcholine transporter, synaptic vesicular amine transporter, vesicular glutamate transporter, excitatory amino acid transporter, dopamine transporter, sodium-dependent serotonin transporter, glycine transporter, but no GABA transporters. Over half of these genes were found to be upregulated in the rediae (Fig. 3B). For a more detailed list of homologues, please refer to Suppl. Table 2 and Supplement information file and Supplement Figure 1A.

#### Vesicle trafficking, release and recycling

We have identified proteins from all core groups ^29^ known for vesicle trafficking, release, and recycling, including 10 synaptotagmins that act as calcium sensors ^30^, Rab proteins important for docking and tethering vesicles, proteins of the SNARE complex, synapsin, transmembrane adenosine triphosphatases, and other proteins. A detailed list of homologues and their domain structures can be found in Supplementary Table 3,4, the Supplementary Information file, and Supplementary Figure 1A.

However, we did not find two chains of synuclein, a multifunctional protein that works as a chaperone in the SNARE complex, Rab GTPase 3A genes, G-proteins that guide synaptic vesicles to active synaptic zones, or Rab effector Noc2, amphiphysin, parvalbumin alpha, and adhesion G protein-coupled receptors.

Of the 75 proteins characterized in this subset, only four were significantly upregulated in cercariae: co-chaperones HSPA8 and PACSIN adapter protein, SNARE complex protein SNAPA, and zinc transporter SLC30A3. Another 26 differentially expressed genes were upregulated in rediae, including two synaptotagmins (SYT2 and -3), which are rediae-stage specific.

#### Reception machinery

Based on gene ontology annotation, over 150 putative G protein-coupled receptors (GPCRs) belonging to all major GPCR groups, and around 100 putative ligand-gated ion channels from three major groups, namely Cys-loop family, glutamate-activated cation channels, and ATP-gated ion channels ^2^, are present in the transcriptome. A selected set of putative receptor proteins were characterized based on reference genes with proved function, including ionotropic and metabotropic receptors, P2X purinoreceptor, soluble guanylate cyclase subunits, some proteins engaged in the second-messenger cascade, and proteins regulating receptor functions.

From the neurotransmitter ligand-gated ion channels, we identified nine nicotinic acetylcholine receptors, orthologs of the acetylcholine gated chloride channels of *Schistosoma mansoni* ACC-1 and ACC-2, inhibitory chloride ion channel glycine receptors, glutamate receptors - cation channels kainite and N-methyl-D-aspartate (NMDA), and chloride channel. From metabotropic receptors, we identified orthologs of muscarinic acetylcholine receptors, dopamine receptors, serotonin receptors, octopamine-tyramine receptors, and orthologs of metabotropic glutamate receptor. We also classified two orthologs of P2X purinoreceptors and nitric oxide receptors soluble guanylate cyclases. A detailed list of homologues, their domain structures and expression profiles can be found in Supplementary Table 5, the Supplementary Information file, and Supplementary Figure 2A.

All the reception machinery genes were expressed at relatively low levels (Supplementary Table 5, Supplementary Figure 2A). Transcripts of a few genes showed equal abundance in rediae and cercariae, including two acetylcholine gated chloride channels, NMDA receptor, metabotropic glutamate receptor, putative metabotropic receptors, putative biogenic amines receptors. Putative serotonin receptor HTR1A2, putative GPCR1 receptor, and soluble guanylate cyclase GUCY1B1 were upregulated in free-swimming larvae. All other 29 DEGs of 48 characterized proteins were upregulated in rediae (Figure 3 B).

We failed to find GABA receptors in neither *C. lingua* nor other Plathelminthes, however, structurally similar multi-pass membrane proteins, the glycine receptors, (GLRB, GLRA) were identified in *C. lingua*. We also did not find homologs of serotonin 5HT3 (Ionotropic), HTR2, 5, 6, 7 receptors and glutamate quisqualate (AMDA) receptors, adrenergic receptors and P2Y receptors in our data set.

We also characterized the proteins involved in electrical synaptic transmission (innexins), proteins maintaining the special electrical properties of the neurons (voltage gated and some other ion channels) and proteins involved in sensory perception.

#### Ion channels

We did not find sodium voltage gated ion channels in the transcriptome of *C. lingua,* only a sodium leak channel non-selective protein (NALCN) that is orthologous to the voltage-independent, cation-nonselective NALCN found in mammals. This protein is permeable to sodium, potassium, and calcium ions (UniProt). Interestingly, the genomes of cestodes and free-living flatworms contain orthologs of the α- subunits forming the core of the voltage-gated sodium channels (SCN1A - SCN11A), while trematodes do not (BLASTp and tBLASTn searches in WormBase Parasite v. WBPS14). We also characterized potassium voltage-gated channels, which are the most diverse group of the ion channel family. Our analysis included representatives of subfamily A, H, Shal, Shab KQT, TWIK, voltage-dependent calcium channels, SK, BK, and homologs of cGMP-gated cation channels. The domain structures and expression profiles are presented in Supplementary Table 6 and Supplementary Figure 2C. In *Sch. mansoni* genome 41 putative channels were identified ^2^ but only few are characterized functionally ^31^. The expression of voltage-gated ion channels in *C. lingua* was relatively low, and all differentially expressed genes were up-regulated in rediae (see Figure 3 B).

#### Innexins

In the transcriptome, innexins are represented by 16 genes (Supplementary Tables 7, the Supplementary Information file, and Supplementary Figure 2B). Only two of these genes are expressed at levels above 100 TMMs, while seven innexins are equally expressed between the studied stages. All differentially expressed innexins were upregulated in rediae (Fig. 3 B).

#### Sensory perception proteins

Proteins involved in sensory perception play a critical role in converting input stimuli into membrane potential changes in sensory neurons. Our research has identified several orthologs, including transient receptor ion channels TRPA1-A18, a non-specific cation channel Piezo, and mechanosensory protein 2 mec-2. Additionally, we have identified several orthologs of light-absorbing opsins, specifically OPN1-4. Supplementary Table 8, Supplementary Information file, and Supplementary Figure 2C provide detailed domain structures and expression profiles.

Among the four opsins identified in the *C. lingua* transcriptome, OPN1 displays a conservative motif indicative of Gq-opsin families. Moreover, two highly conserved peptide amino acids histidine and proline in the carboxy terminal intra-cellular loop domain show significant (64-67%) identity to the characterized rhabdomeric (r) Gq-opsins of *Sch. haematobium* and *Sch. mansoni* (Smp_104210 and Sha_101185) ) ^32^, which have a suggested role in photokinesis ^33^. Rhabdomeric opsins are known to attain color vision in invertebrates, with higher sensitivity to blue light ^34^. Interestingly, the expression of *Cl*OPN1 is higher than other putative opsins (Supplementary Figure 2C) in both studied stages, which corresponds to the experimental data on the predominant blue-light sensitivity of *C. lingua* cercaria ^8^. Most of the differentially expressed genes (DEGs) in this subgroup were upregulated in cercariae developing in rediae, while two TRPA channels (*Cl*TRPA10 and 16) were upregulated in freely swimming larvae (Figure 3B).

We did not find degenerins, mechanosensory abnormality protein 6, transmembrane channel-like proteins, mechanosensory transduction channel NOMPC, and amiloride-sensitive sodium channel subunits in the transcriptome of *C. lingua*.

In our study, we characterized 229 neuronal transcripts from the *de novo* transcriptome of *C. lingua*. We confirmed the existence of most neurotransmitter systems and key proteins involved in the neurotransmission machinery, however a few were found to be absent (Figure 3A). The majority of differentially expressed genes (DEGs) were upregulated in rediae containing all stages of cercariae development.

### The HCR *in situ* hybridization of selected neuronal genes has revealed their expression in the nervous system

Although most of the neuronal proteins were upregulated in rediae samples, only a few were upregulated in cercariae. We hypothesized that the dispersal larva strategy is focused on saving energy at all costs, and therefore the neuronal proteins upregulated in non-feeding larvae should play a crucial role in dispersal swimming, host finding, or regulation of metabolism. The localization of expression of certain genes can provide clues about their function, which is why we performed HCR *in situ* hybridization of selected transcripts upregulated in dispersal larvae.

We proved that characterized neuronal genes are expressed in the nervous system of *C. lingua*. To do this, we used staining with neuronal marker antibodies (AB) and HCR *in situ* hybridization for the same transcripts in parallel. We were aware of the limitations of using commercial AB, which were not previously tested on trematode proteins. Nevertheless, we managed to compare the localization of AB staining with the results of *in situ* HCR based on the anatomy of the nervous system described earlier ^35, 36^. All three neuronal markers used: calcium sensor synaptotagmin, ubiquitous in the nervous system ^37^; CHAT, enzyme essential for production of acetylcholine, and serotonin or SERT – specific serotonin transporter; were localized or stained in the nervous system. They were localized both in the central nervous system, cerebral ganglia, commissure, and ventral nerve cords, as well as in peripheral neurons, nerve cords, and nerves of cercaria. Interestingly in contrast to AB staining, no neuronal markers expression was localized in the nervous system of redia itself, only in developing cercaria within redia (Fig 4.).

**Figure 4.**
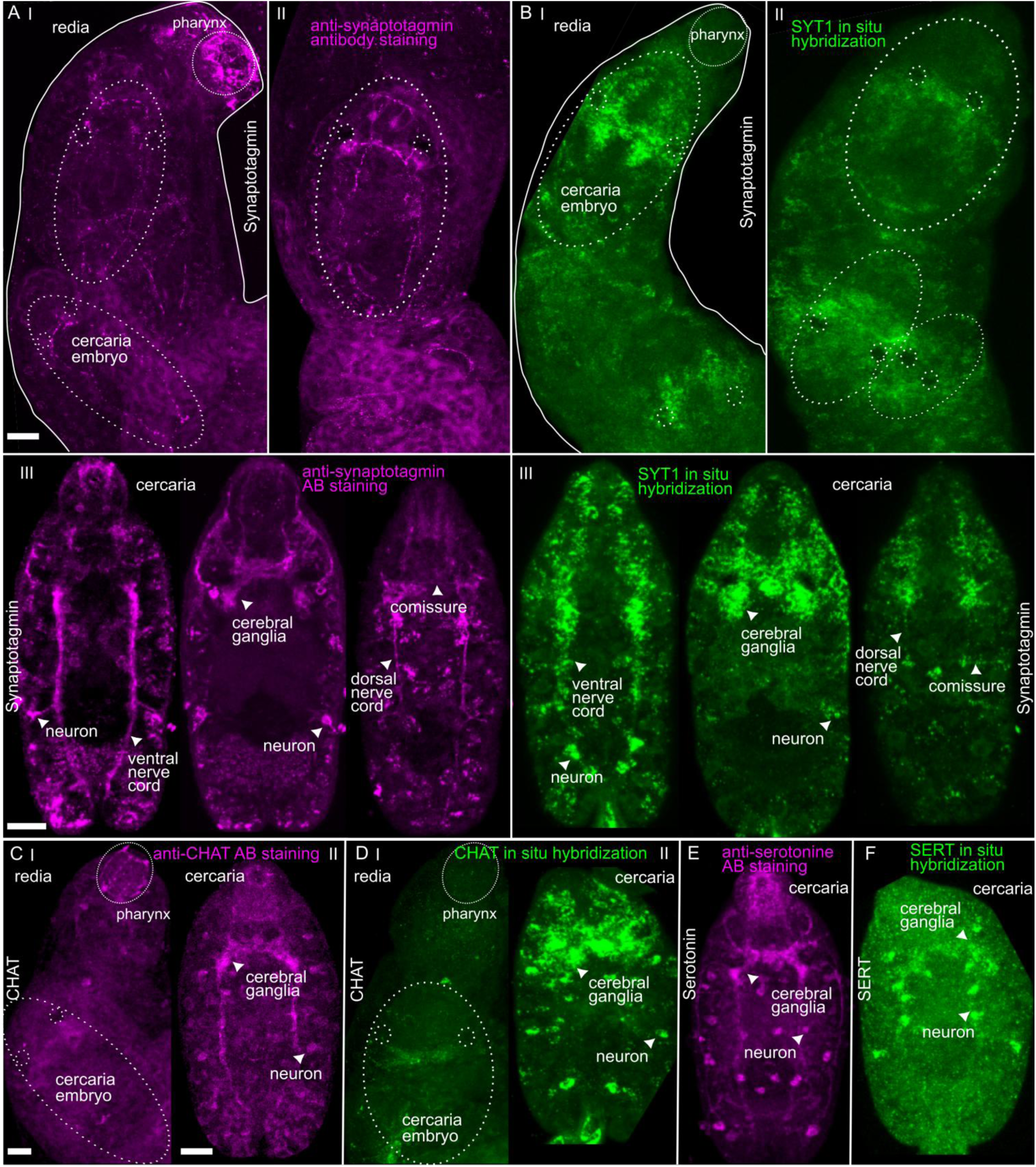
Parallel staining of neuronal markers by antibody staining and HCR *in situ* hybridization confirmed their localization in the nervous system. (**A**) Antibodies staining against SYT1 (magenta) and (**B**) HCR *in situ* hybridization of *Cl*SYT1 gene (green) in the nervous system of redia and developing cercaria inside (marked with dash circles) in upper (**I**) and lower part of redia (**II**) and optical sections of mature cercaria removed from the redia (**III**). (**C**) Antibodies staining against CHAT (magenta) and (**D**) HCR *in situ* hybridization of *Cl*CHAT gene (green) in the nervous system of redia and developing cercaria inside (marked with dash circles) (**I**) and in mature cercaria removed from the redia (**II**). (**E**) Antibodies staining against serotonin (magenta) and (**F**) HCR *in situ* hybridization of *Cl*SLC6A4 (SERT) gene (green) in the nervous system of mature cercaria removed from the redia. Note signed parts of the central nervous system and peripheral neurons. Scale bar = 50 µm

### Expression of genes upregulated in dispersal larva suggests their role in the nervous system maintenance and facilitating host finding

The use of HCR *in situ* hybridization allowed for the quantification of mRNA levels ^38^ for three selected genes, namely *Cl*SYT1, *Cl*CHAT, and *Cl*SNAP, providing additional evidence for our DEG analysis. We compared the fluorescence in the central nervous system between three samples, redia, developing cercaria, and freely swimming cercaria (Fig 5A) and found the localization of neuronal markers in the nervous system of developing and freely swimming cercaria but not in redia. The results of HCR quantification were generally consistent with our RNAseq data (Fig 5 B, C, D).

**Figure 5.**
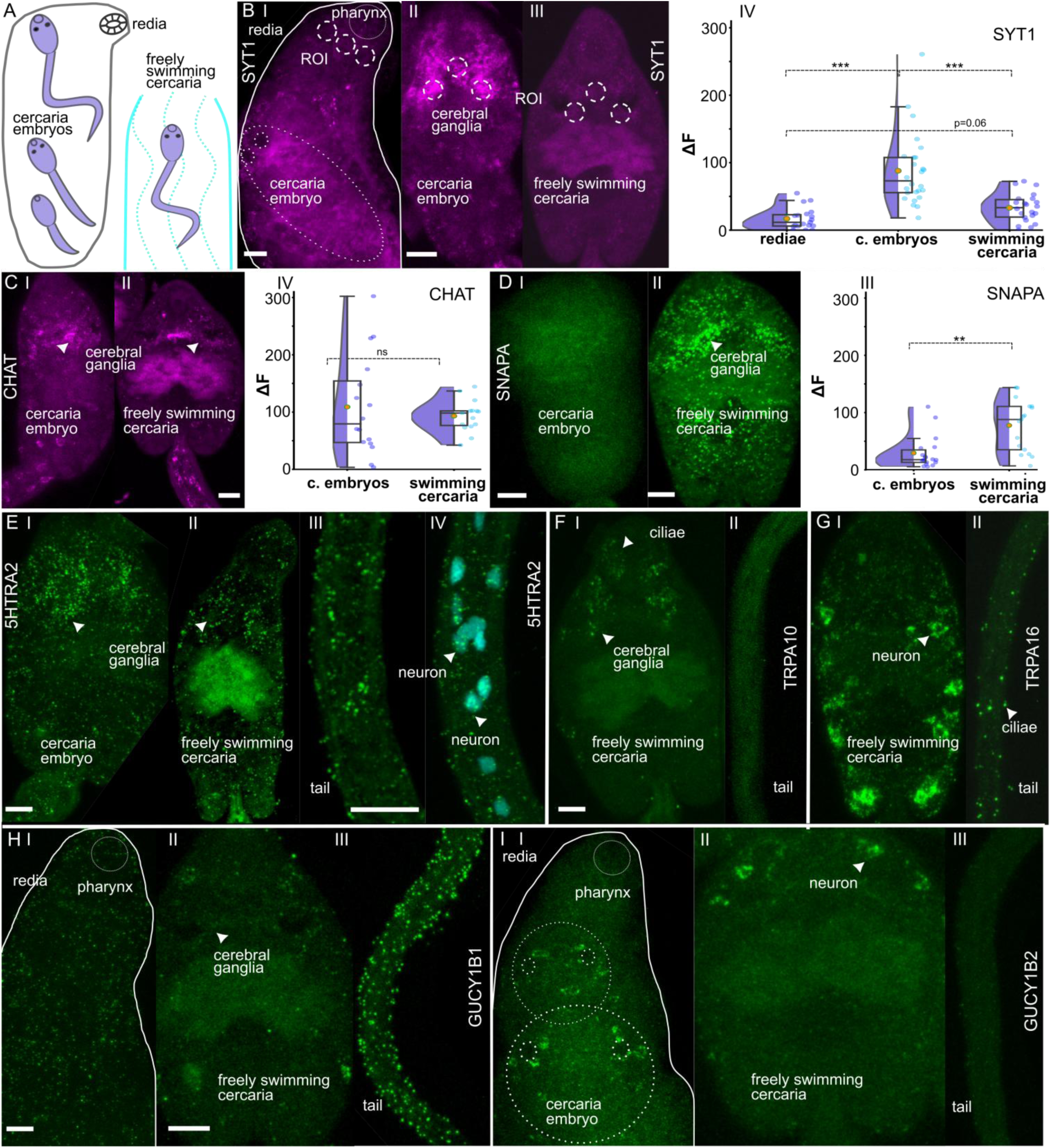
Localization of dispersal “stage-specific” genes in cercaria highlight their potential function. (A) Stages studied (redia with developing cercaria and freely swimming cercaria). (**B - C**) Expression of neuronal markers (magenta) between stages and quantification of HCR fluorescence. (**B**) Synaptotagmin expression in redia (**I**), cercaria embryo (**II**) and freely swimming cercaria (**III**) and its quantification (**IV**). (**C**) CHAT expression in cercaria embryo (**I**) and freely swimming cercaria (**II**) and its quantification (**III**). (**D_I**) Expression of DEGs upregulated in dispersal cercaria stage (green). (**D**) SNAPA expression in cercaria embryo (**I**), and freely swimming cercaria (**II**) and its quantification (**III**). (**E**) 5HTRA2 gene expression in cercaria embryo (**I**), and freely swimming cercaria body (**II**), its tail (**III),** and with nuclei stained with DAPI **(IV**). (**F**) TRPA10 gene expression in freely swimming cercaria body (**I**), and tail (**II).** (**G**) TRPA16 gene expression in freely swimming cercaria body (**I**), and tail (**II).** (**H**) GUCYB1 gene expression in redia (**I**) but not in cercaria embryos, freely swimming cercaria body (**II**), and tail (**III).** (**I**) GUCYB2 gene expression in redia and in cercaria embryos (**I**), freely swimming cercaria body (**II**), and no expression in the tail (**III).** Note signed parts of the central nervous system and peripheral neurons. Scale bar = 50 µm. Test = Mann–Whitney U test or Kruskal-Wallis analysis of variance with the Dunn‟s posthoc test. P values: * <0.05; ** <0.01; *** <0.001

To further investigate the expression of a set of neuronal genes upregulated in dispersal larvae, we localized the transcripts of selected upregulated genes. Alpha-soluble NSF attachment protein *Cl*SNAPA plays a role in the fusion of synaptic vesicles to the plasma membrane ^39^, while *Cl*PACSIN1 is involved in synaptic vesicle endocytosis ^40^. We assumed that both transcripts should be localized in synapses, and as a result, expression was found to be distributed along the body of the cercaria with an aggregation in the central nervous system (Fig 5 D, Supplement Figure 3). Putative serotonin receptor *Cl*HTR1A2, on the other hand, was detected in the cerebral ganglia and in the neurons of the tail (Fig 5 E).

The transcripts of the receptor-activated non-selective cation channels *Cl*TRPA10 and *Cl*TRPA16, homologs of polymodal TRPA1 with suggested temperature, chemical and mechanosensitivity ^41^, were found to be expressed differently within the aggregation of cilia in the area around and below the oral sucker for TRPA10 (Fig 5 F), as well as at the surface of the tail and in the set of peripheral neurons on both sides of the body for TRPA16 (Fig 5 G). Additionally, the soluble guanylate cyclases ClGUCY1B1 and ClGUCY1B2, homologs of the nitric oxide receptor GUCY1 protein ^42^, were found to be expressed differently. While GUCY1B1 had low expression on the surface of redia and not visible expression in developing cercariae, it had low expression in the cerebral ganglia and strong surface expression in the tails, the main organ of locomotion in the freely swimming larvae (Fig. 5 H). GUCY1B2, on the other hand, was expressed in two symmetrical pairs of neurons in the apical part of the body in developing cercaria inside redia and in the freely swimming larvae, but not in redia tissues (Fig 5 I).

## Discussion

### Differentially expressed genes

The isolation of parthenitae (redia) stage tissues for independent analysis is challenging as rediae contain developing cercariae embryos that cannot be separated from maternal body. Therefore, our study’s "redia" samples are mixed and predominantly contain cercarial tissues at all stages of development, from early embryos to full-grown cercariae ready to shed (Fig 1AII, Fig 4, 5). We assume that some of neuronal genes expressed in the grown-up larvae inside rediae can be utilized in complex and uncharacterized navigation and motility inside the snail host and during the shedding.

In our dataset, we observed that 14.5% of overall protein-coded genes were upregulated in rediae, while only 3.2% were upregulated in freely swimming cercariae. Similar stage-specific differences have been reported recently in other parasitic flatworms, such as *Fasciola gigantica* ^43^ and two species of psilostomatids ^24^. Our GO analysis results partially support previous findings on model species *Schistosoma mansoni*, where cercariae were described as a "silent" stage with inactive cell division and upregulated genes involved in energy metabolism ^44, 45^. This similarity could be explained by the similar swimming strategy of *Sch. mansoni* and *C. lingua*, where both show intermittent swimming with active bursts and sinking phases believed to increase their life span and probability of infecting the next host ^5^. For the identified neural genes 103 genes or 45.6% were upregulated in redia compared to 11 or 4.8% upregulated in dispersal cercaria. While a few upregulated genes of certain neuronal families (GPCRs, Ion channels) were observed in cercaria of *Sch. mansoni* ^44^, these did not match the list of upregulated genes in dispersal cercaria of *C. lingua*. However, a large proportion of genes were constantly expressed between stages in *C. lingua*, and differences in the composition of upregulated genes raise questions about species-specific gene expression profiles related to differences in behavior cues and strategies for infecting the next host. These cues and strategies differ between mammal-infecting *Sch. mansoni* and fish-infecting *C. lingua*.

We believe that the dispersal larvae of trematodes exhibit a highly specialized functionality, primarily focused on energy store digestion (metabolism), motor coordination, and navigation. During the resource-rich feeding stage of the rediae, complex neuronal proteins, including ion channels, are synthesized, indicating the crucial role of this stage in the development of the nervous system in cercaria embryos.

Our data on the expression of neuronal genes upregulated in dispersal cercariae included a few genes involved in synaptic vesicle trafficking and release. We found two TRPA channels that might be related to sensation, a putative GPCR receptor and two putative NO receptor soluble guanylate cyclase. A recent study showed active transcription in cercariae of *Schistosoma mansoni* localized in the tail region, including vesicle transport GO transcripts ^46^. In contrast, in our data expression of genes *Cl*SNAPA and *Cl*PACSIN1, involved in synaptic vesicle fusion and endocytosis, and GPCR *Cl*HTR1A2 was localized in the central nervous system and all over the body and tail of cercaria. For more details on TRPAs and SoGUCs see below.

### Classical neurotransmitters pathways

Our research has revealed the absence of gamma-amino butyric acid (GABA) and histamine signaling pathways, as well as epinephrine and octopamine biosynthesis in *C. lingua*. These findings are consistent with previously published data based on genome analysis of few parasitic flatworms ^47, 48^, which have been discussed before ^1, 49, 50^. Interestingly, despite the lack of genes for histamine biosynthesis and metabolism, one of the upregulated GPCR receptors in cercaria (*Cl*GPCR1) is similar to the *Sch. mansoni* GPCR (with 29% similarity) that has been shown to have affinity for histamine ^51^. Based on our transcriptome analysis, some special features of classical neurotransmitter pathways in *C. lingua* were observed. These include the similarity of ClSLC6A2 to both the Sodium-dependent dopamine transporter and the Sodium-dependent norepinephrine transporter, as well as the chloride channel nature of ionotropic acetylcholine receptors, as previously described in *Sch. mansoni* ^50, 52–55^. Moreover, the homologs of neuronal proteins identified in *C. lingua* are also similar to the genes involved in classical neurotransmitter pathways that have been functionally characterized in other trematode species ^49, 56–60^.

### The loss of NOS may have shaped the complex life cycle of trematodes

Among the small gaseous messenger pathways in the nervous system, we found only orthologs involved in hydrogen sulfide biosynthesis in *C. lingua*. We discovered that *C. lingua* and 33 Platyhelminthes species, representatives of classes Trematoda, Monogenea, Cestoda, and free-living turbellarians *Macrostomum lignano* and *Schmidtea mediterranea* have lost the nitric oxide synthase (NOS) gene, which is essential for the cellular signaling NOS pathway ^61^. We believe that two histochemical methods used to mark the presence of NOS in the tissues of parasitic flatworms based on antibody staining ^62^ and NADPH diaphorase histochemical staining ^63^ might have marked unspecific targets, as described earlier ^64^. However, it is important to note that the genes encoding NOS exist in other lophotrochozoans ^65, 66^ (Figure 6). We propose that the loss of nitric oxide synthase in Platyhelminthes was compensated by the peculiarities of the parasitic lifestyle of these animals, which allows for unrestricted use of host-derived NO. Similar losses of the gene for this enzyme were reported for representatives of Ctenophora ^67^ and for nematodes in which it has been attributed to consequences of life in an environment enriched with nitric oxide by bacteria ^68^. However, the lack of NOS in trematodes does not mean the absence of the NO signaling pathway. The upregulation of soluble guanylate cyclase in the *C. lingua* cercaria may indicate the involvement of NO in energy metabolism, motility, or adaptation to lower concentrations of non-enzymatically produced NO ^69^ in seawater compared to host tissues. And in fact, we localized the expression of two studied soluble guanylate cyclases in both tail, the main organ of locomotion, and body neurons suggests its crucial involvement in the dispersal process and host detection (sensation). Nevertheless, the source of NO in cercariae outside the host body remains enigmatic and needs further study. Moreover, it was shown that bacterially derived NO enhances *C. elegans* longevity and stress resistance ^68^. We speculate that NO-deficiency may be a similar or even more significant reason for the short lifespan ^8, 70^ of cercariae than energy limitations. It may have also played a crucial role in the evolution of digenean homo-, di-, and trixenic life cycles, where dropout of free-swimming larvae is common ^23^. Additionally, the loss of NOS genes in both Platyhelminthes and Nematoda ancestors (Figure 6) may have prompted the shift from a free-living to a parasitic lifestyle. To test this hypothesis, further studies are necessary.

**Figure 6.**
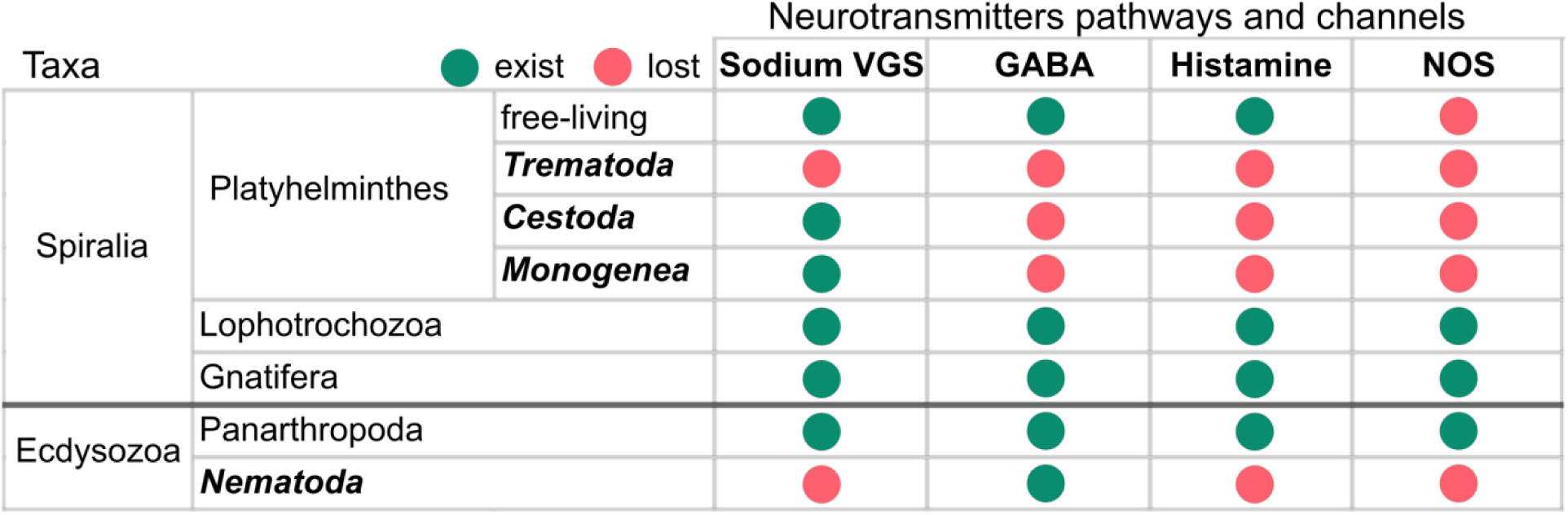
The profiles of neuronal gene losses observed in Platyhelminthes exhibit parallels with those found in Nematoda across various groups within the Protostomia. The taxa with the evolutionary transition from free-living to parasitic lifestyle are marked in bold.

### Synaptic transmission

The genes composition involved in synaptic vesicle cycle in bilaterians is conservative and so as genes composition of synaptic vesicle cycle in *C. lingua*. Our data has enabled us to identify transcripts that encode all core components of animal neurosecretory vesicles and machinery of synaptic vesicle cycle: V-ATPases, vesicular neurotransmitter transporters, transporter and transporter-like proteins, proteins with four transmembrane domains, synapsins, synaptotagmins, secretory SNAREs, endosomal SNAREs, transiently associated proteins SNARE binding partners and co-chaperones ^26, 29, 65^ which is corresponding to recent identification of synaptic transmission machinery in different life cycle stage juveniles of liver fluke *Fasciola hepatica* ^3^.

### Electric properties of the neurons

Neurons use the potential energy in transmembrane ionic potentials regulated by ion channels to drive regenerative traveling electrical signals called action potentials (AP) that carry signals down axons ^71^. APs are a widespread signaling phenomenon in living organisms and in animals the fast AP is usually sodium based. Sodium voltage gated channels (SVGC) thought to have evolved alongside the early neuromuscular systems being evolutionary related to calcium voltage gated channels ^71, 72^. The absence of sodium voltage gated channels as in *C. lingua* was also described in *Sch. mansoni* ^2^ but not in other parasitic (Cestoda, Monogenea) and free-living flatworms clades ^50, 73–75^. Interestingly, we recorded APs in the tail of the *C. lingua* cercariae earlier ^76^. The occurrence of APs in the absence of SVGC is known for nematodes and explained by the involvement of potassium and calcium VGC ^77^, that can be the case in *C. lingua* APs generation.

### Sensory perception of cercaria

Cercaria of *C. lingua* has a complex behavior with characterized light and water turbulence perception playing a key role in finding the fish host ^7, 8, 21, 78^. The presense of rhabdomeric (r) Gq-opsins in *C. lingua,* as well as in of *Sch. haematobium* and *Sch. mansoni* ^32^ along with previous reports of selectivity to different light spectra ^8^ suggests promising avenues for future experiments. Certain transient receptor potential ion (TRP) channels are associated with r-opsins and necessary for phototransduction ^34^. Another TRPs can be involved in mechanotransduction ^79^ or thermosensation ^41, 80^. We suggested that at least some of TRP channels identified play a role in water turbulence perception and finding the fish host ^81^. Cercariae *C. lingua* have multiple cilia at their surface ^82^ some of the long ciliae shown to have connections to neurons ^83^. In fact, two TRPA channels upregulated in cercaria were primarily expressed in the short cilia and in a set of peripheral neurons, highlighting their potential importance in the process of mechanosensation/transduction or heat/cold sensing ability and a next host finding.

### Evolutionary perspective

The loss of neurotransmitter pathways is a prevalent occurrence observed across diverse taxa^65^. Within the Protostomia, the convergence between nematodes and trematodes highlights their shared absence of specific neurotransmitters and sodium voltage-gated channels (Figure 6), which may imply a loss either before or after their specialization as parasites, or potentially a combination of both scenarios. Remarkably, the absence of nitric oxide synthases (NOS) in all taxa of Platyhelminthes, including both free-living and parasitic forms, strongly indicates the absence of NOS during an earlier phase in the evolutionary history of flatworms. It suggests a plausible evolutionary trajectory wherein the absence of particular neurotransmitter systems predates the specialization of proto-worms as parasites and can have potential relationship with the complex life cycles and other parasitic adaptations. Further investigation is necessary to understand how this combination of neurotransmitter systems interact, modulate neural activity, and contribute to the overall functionality of the nervous system.

## Conclusions

This article outlines the first survey of neuronal genes/proteins identified, characterized and validated for a novel model species for trematode biology, *Cryptocotyle lingua*. This parasite demonstrates the surprising absence of certain key signaling pathways and ion channels. We further demonstrate that specific molecular building blocks of its nervous system exhibit differences in molecular pathways and gene expression between the different life stages, i.e. the redia and cercariae stages.

Several groups of neuronal proteins of trematodes, such as various channels and receptors, have previously been identified as potential targets for anti-parasitic treatment ^54, 84, 85^. However, further research is required to understand their mechanisms of action to develop effective treatments via these targets. Additionally, the development of resistance to these treatments may limit their long-term efficacy, highlighting the need for continued research into new therapeutic targets. To this end, having a model system such as *C. lingua*, which has an easily quantifiable behavior and a molecularly accessible and characterized nervous system, could prove beneficial for drug screenings and research in this field.

## Methods

### Animal samples

Cercariae and rediae of *Cryptocotyle lingua* (Creplin, 1825) (Digenea: Heterophyidae) were obtained from naturally infected common periwinkles *Littorina littorea* (Linnaeus, 1758), which were collected at the Biological station of Zoological Institute RAS “Kartesh” (Chupa inlet, Kandalaksha bay, the White Sea) and in the vicinity of Bergen (Tyssøyna, the North Sea). To collect the cercariae, each infected mollusk was placed individually in a reservoir filled with sea water of natural salinity (24‰). After four hours of cercariae emission, the larvae were collected and identified ^86^ under a dissection microscope. In order to clear the larvae sample of the mollusk mucus and other large xenogenic contaminations, cercariae were placed in a dark side of sterile 100 mm Petri dish filled with sterile seawater (SSW: 22 μm filtered) where they migrated to opposite side of the dish due to positive phototaxis. The cercaria were accumulated in 10 ml sterile tubes and cooled on thawing ice to 4°C, which allowed the larvae to lose their mobility to be concentrated at the bottom of the tube. The original water was removed, and the larvae were rinsed with SSW in four passages. Each sample of living cercariae contained 600-800 individuals.

The rediae were recovered from the same mollusks after their dissection under a microscope and collected in a sterile Petri dish filled with SSW. Harvested rediae were immediately transferred to sterile round-bottom plastic wells for washing from scraps of host tissue and mucus. The washing with SSW was controlled with thin pipet, carefully removing the remnants of host-derived tissues, cell aggregates, injured rediae and cercariae released during the process. In general, each sample of live rediae (400-500 individuals) was washed 15 times in a total volume of 150 ml of SSW. To prevent adhesion of both rediae and cercariae to the surface of a sterile hydrophobic plastic, it was treated with a sterile solution of bovine serum albumin (BSA: 0.01 mg/ml). A part of the infected snail was kept in the aquarium facility of the Michael Sars Center, University of Bergen and was used for fixations and behavior experiments.

### RNA isolation and libraries preparation

Total RNA isolation was performed using ExtractRNA™ (Evrogene, Moscow, Russia) according to the manufacturer’s instructions, followed by intensive pipetting to fully lyse rediae and cercariae tissues. The RNA samples were stored in 80% EtOH at -80°C for no longer than two weeks. Library construction for transcriptome sequencing was carried out using the TruSeq Stranded mRNA LT Sample Prep Kit from Illumina (San Diego, CA), following the manufacturer’s protocol. We obtained three paired biological replicates for two life cycle stages: rediae and cercariae from the same infected periwinkle in one pair. Sequencing of the six libraries was performed at Macrogen (Seoul, Korea) on an Illumina HiSeq 4000 platform with paired-end 2 × 100 reads.

### Species identification

Morphological identification of species in digeneans can be challenging, even when using adult worms, and is sometimes impossible when using their larval stages ^24, 87^. To address this issue, we searched the transcriptome of *C. lingua* for the sequences of i) internal transcribed spacers 1 and 2 (ITS1 and ITS2), located between 18S and 28S in ribosomal RNA (rRNA), and ii) the mitochondrial cytochrome c oxidase subunit 1 gene (COX1). We found both complete sequences encoding 18S-ITS1-5.8S-ITS2-28S rRNA (GenBank: MW361240) and COX1 (GenBank: MW361241) to be 100% identical to the previously published sequences of *C. lingua* ^88^. Thus, our morphological identification of the species, and the absence of cryptic species, is supported by molecular identification.

### *De novo* assembly and differential expression

The quality of paired-end read data was assessed using FastQC (v0.11.9). Adapter and low quality bases (Phred score below 30) were trimmed from sequence reads with Trimmomatic (v0.38) ^89^. Sequences longer than 50 bp were retained for further analyses. The prepared paired-end read data from six libraries were pooled together and used in *de novo* assembly of the reference transcriptome using Trinity (v2.8.6) ^90^ with strand-specific read orientation and a minimum contig length of 200 bp. This assembly was further reduced by removing the low-expressed (FPKM<3) and several xenogenic transcripts.

A local BLASTn approach was used to filter out potential xenogenic contaminating sequences from the raw assembly: with the host-mollusk *Littorina littorea* transcriptome of hemocytes and kidney (NCBI TSA: GGCG00000000.1) ^91^, whole body transcriptomes of related species *L. obtusata, L. fabalis, L. saxatilis*^92^ and human genome (Encode: GRCh38, primary assembly). SILVA rRNA database ^93^ was also used to search other potential xenogenic contaminations. The quality of the curated assembly was assessed by the presence of the Metazoa single-copy orthologues and verified using BUSCO (v4.1.4; metazoan-odb10) ^94^. Only the longest proteins were selected as representatives of each gene for BUSCO-benchmarking.

RNA-seq analysis was performed with Bowtie2 (v2.3.5.1) ^95^ and quantification method RSEM, to obtain gene expression values including counts, TPM (Transcripts Per Million) and TMM (TMM-normalized TPM matrix). To identify differentially expressed genes (DEGs) in rediae and cercariae, an EdgeR test ^96^ for paired samples was carried out with the reference transcriptome and stage specific reads in three replicates. Transcripts with absolute fold change (FC) values ≥4 and a false discovery rate (FDR)-corrected p-value ≤0.05 were considered as differentially expressed. To avoid the loss of biologically significant differences, an additional detailed analysis for verified neuronal genes was performed using a two-tailed T-test for paired TMM data that passed the Shapiro-Wilk normality test.

### Aminoacid sequences prediction and functional annotation

The coding regions within transcript sequences were identified using the TransDecoder tool with either BLASTp (Swissprot) and PFAM hits as the ORF retention criteria. The predicted proteins were then annotated using the Trinotate tool with HMMER/PFAM protein domain identification, signalP (v4.1), TMHMM (v2.0c) and BLASTp (e-value threshold 1e-25) against the UniProt/SwissProt protein database. Additionally, protein families were annotated using InterProScan, and gene ontology (GO) mapping (e-value cut-off 1e-25) was conducted in Blast2GO, including the curated transcriptome as the reference set. An enrichment gene ontology (GO) analysis for the protein coding DEGs (test set) was also performed in Blast2GO, and a Fisher‟s exact test was run with an FDR cut-off of 0.05. Only over-represented biological processes (BPs) were further analyzed.

### Identification of neuronal genes

We conducted a similarity analysis and protein domain architecture study on neuronal genes, using reference proteins involved in neurotransmission, sensory and neural pathways obtained from the Uniprot database. We primarily selected *Homo sapiens* proteins as reference species, but in cases where functionally characterized homologs were available, or if the reference proteins were absent in vertebrates, we used proteins from the trematode species *Schistosoma mansoni* or another invertebrate model species. To identify candidate homologs, we locally tBLASTn searched the *C. lingua* transcriptome with an E-value threshold of 1e-5 for short sequences. Any hits below this threshold were considered as candidate homologs. We examined all candidate protein sequences for the presence of conserved amino-acid motifs related to Pfam ^97^ and SMART ^98^ domains. We used the InterProScan tool in Geneious R10 (https://www.geneious.com) to analyze protein domain architectures. We considered proteins whose domain content and order were identical to the corresponding prototypes as homologous. Supplementary Tables 1 – 7 contain information on the reference genes, E-value of similarity, and NCBI accession numbers of homologous *C. lingua* transcripts.

### Hybridization chain reaction (HCR) *in situ* hybridization

Probes sets for HCR *in situ* hybridization were generated by insitu_probe_generator ^99^ and produced by Integrated DNA Technologies, Inc. (USA) for the following gene models: ClCHAT, ClSYT1, ClSLC6A4 (SERT), ClSNAP, ClPACSIN1, ClHTR1A2, ClGUCY1B2, ClGUCY1B1, ClTRPA16, ClTRPA10, as well as for sense control sequences for ClCHAT and ClGUCY1B2. From 27 to 33 split initiator pairs were used for each gene (Supplement Table 9).

Embryos were fixed in 4% formaldehyde overnight at 4°C. Samples were then washed two times in phosphate-buffered saline and 0.1% Tween 20 (PBST). Samples were then dehydrated in 50% methanol and then with 100% methanol, and stored in 100% methanol at −20°C until use.

The HCR protocol was based on a previously published protocol for fruit fly larva ^38^. For a control experiment, we used negative control sense sequences and HCR reactions without probes. After HCR animals were stained with DAPI and mounted on slides in 50% glycerol. All animals for HCR *in situ* hybridization were stained at once using the same hybridization, washing and amplification time for all genes.

### Antibody staining

For antibody staining, animals were fixed in 4% formaldehyde overnight and kept in PBST at 4°C. We used commercial antibodies including anti-serotonin rabbit, anti-synaptotagmin rabbit (Sigma), anti-CHAT goat (Invitrogen), Alexa Fluor 488 donkey anti-rabbit, and Alexa Fluor 555 donkey anti-goat (Molecular Probes). Prior to staining, animals were treated with Proteinase K and incubated overnight in blocking solution. We stained animals with primary and secondary antibodies for 5 days at a 1:800 dilution in PBST containing 1% BSA (Sigma). After washing in PBST, animals were stained with DAPI and mounted on slides in 50% glycerol.

### Imaging

HCR *in situ* and antibody stained samples were imaged using Olympus fv3000 and Leica Stellaris 8 microscopes using a 40× objectives. Z-sections of whole animals were taken with the same laser power and settings for all HCR *in situ* animals.

### Quantification of HCR *in situ* signals

For each gene, the mean fluorescence intensity of three identical diameter circular regions of interest (ROI) was measured in ImageJ (NIH) for the two cerebral ganglia and a cerebral commissure at the Z-section of the animal. The background signal was measured for the random region within the animal outside the central nervous system localization and subtracted from the mean fluorescence of three ROIs.

### Behavior recording and video analysis

We used behavior setup and videos analysis described earlier ^100^ using ToxTrac program with default settings ^101^, when the coordinates of the center of mass of each animal in each image frame were identified. We recorded animals for 5 minutes in the absence of visible light and constant 15° C in Artificial sea water (ASW). Later all positional data was calibrated to the size of the arena to obtain movement trajectories and to calculate the speed and path complexity for the animal trajectories as described ^100^.

### Data visualization and statistics

All data analysis and visualization were performed with Python 3.11 using the numpy, pandas, scipy, scikit-learn, matplotlib and seaborn libraries. Samples were checked by Shapiro-Wilk normality test and compared with Kruskal-Wallis analysis of variance with the Dunn‟s posthoc test and Mann–Whitney U tests.

### Availability of data

The data that support the findings of this study are openly available in GenBank and Figshare. The raw data have been deposited in DDBJ/EMBL/GenBank (Bio-Project: PRJNA646866; BioSample: SAMN15567434, SAMN15567352; SRA accession: SRR12246812 - SRR12246817). The curated *C. lingua* Transcriptome Shotgun Assembly (TSA) has been deposited at DDBJ/EMBL/GenBank under the accession GISJ00000000. Both, raw and curated (v1.2.2) assemblies, are available at Figshare (https://doi.org/10.6084/m9.figshare.12673415.v1; https://doi.org/10.6084/m9.figshare.13386671.v1; https://doi.org/10.6084/m9.figshare.13387091.v1).

## Acknowledgements

We are grateful to Dr Alexander Klimovitch and Dr Kirill Nikolaev for their kind help in initial sampling. M.C. and O.T. were supported by two grants of the Research Council of Norway: grant number 339399 to M.C. and grant number 234817 to Sars International Centre for Marine Molecular Biology Research (supporting both M.C. and O.T.). AG is supported by the IEPhB Research Program 075-0152-22-00 and IEPhB Research Resource Center. We thank Mie Wong for comments on the manuscript.

