## Supplement_information_file for "When offspring outsmarting parents: Neuronal genes expression in two generations of marine parasitic worm"

Here we characterize the domain structures and provide expression profiles for the identified neuronal proteins.

*Neurotransmitters biosynthesis and metabolism*

We identified a group of cytoplasmic proteins homologues of enzymes involved in the biosynthesis and metabolism of classical neurotransmitters and small gaseous messengers in the transcriptome of *C. lingua* (Supplement table 1). Biopterin-dependent aromatic amino acid hydroxylases contain biopterin_H catalytic domain that catalyzes the ring hydroxylation of aromatic amino acids and act as rate-limiting catalysts for catecholamines biosynthesis. We identified orthologs of biopterin-dependent aromatic amino acid hydroxylases, tyrosine 3-monooxygenase and tryptophan 5-hydroxylase 1 (*Cl*TH, *Cl*TPH1). Pyridoxal-dependent decarboxylase contain the conserved domain of group II pyridoxal-dependent decarboxylases and catalyses the decarboxylation of tryptophan to tryptamine, tyrosine into tyramine and histidine to histamine. We classified aromatic-L-amino-acid decarboxylase and tyrosine decarboxylase (*Cl*DDC, *Cl*TDC). The copper type II ascorbate-dependent monooxygenase family proteins contain Cu2 monooxygen domains that require copper as a cofactor and which uses ascorbate as an electron donor for monooxygenase activity. We identified dopamine beta-hydroxylase (*Cl*DBH) in *C. lingua* transcriptome. Aldehyde dehydrogenases contain aldehyde dehydrogenase domain and oxidize aliphatic and aromatic aldehydes including norepinephrine aldehyde using NADP as a cofactor. We classified aldehyde dehydrogenase family 3 member A2 (*Cl*ALDH3A2). The choline/carnitine acyltransferase domain is found in a number of eukaryotic acetyltransferases including choline o-acetyltransferase an enzyme that catalyses the biosynthesis of the neurotransmitter acetylcholine. We identified ortholog of choline acetyltransferase choline O-acetyltransferase (*Cl*CHAT) in the transcriptome of *C. lingua*. Carboxylesterase family type B proteins containing the carboxylesterase domain and act on carboxylic esters including acetylcholine. We classified orthologs of type-B carboxylesterase/lipase family acetylcholinesterase and cholinesterase 1,2 (*Cl*ACHE, *Cl*BCHE1, *Cl*BCHE2) involved in the in the transcriptome of *C. lingua*. Glutaminases contain eponymous glutaminase domain and deaminate glutamine to glutamate. We classified glutaminase *Cl*GLS. Cys/Met metabolism PLP (pyridoxal-5'-phosphate)-dependent enzyme family contain eponymous domain and includes enzymes involved in cysteine and methionine metabolism which use PLP as a cofactor. We classified cystathionine gamma-lyase *Cl*CTH. PLP-dependent enzymes superfamily contain a PLP domain and includes cysteine synthase. We identified cystathionine beta-synthase *Cl*CBS in transcriptome of *C. lingua* (Suppl. Figure 1 B).

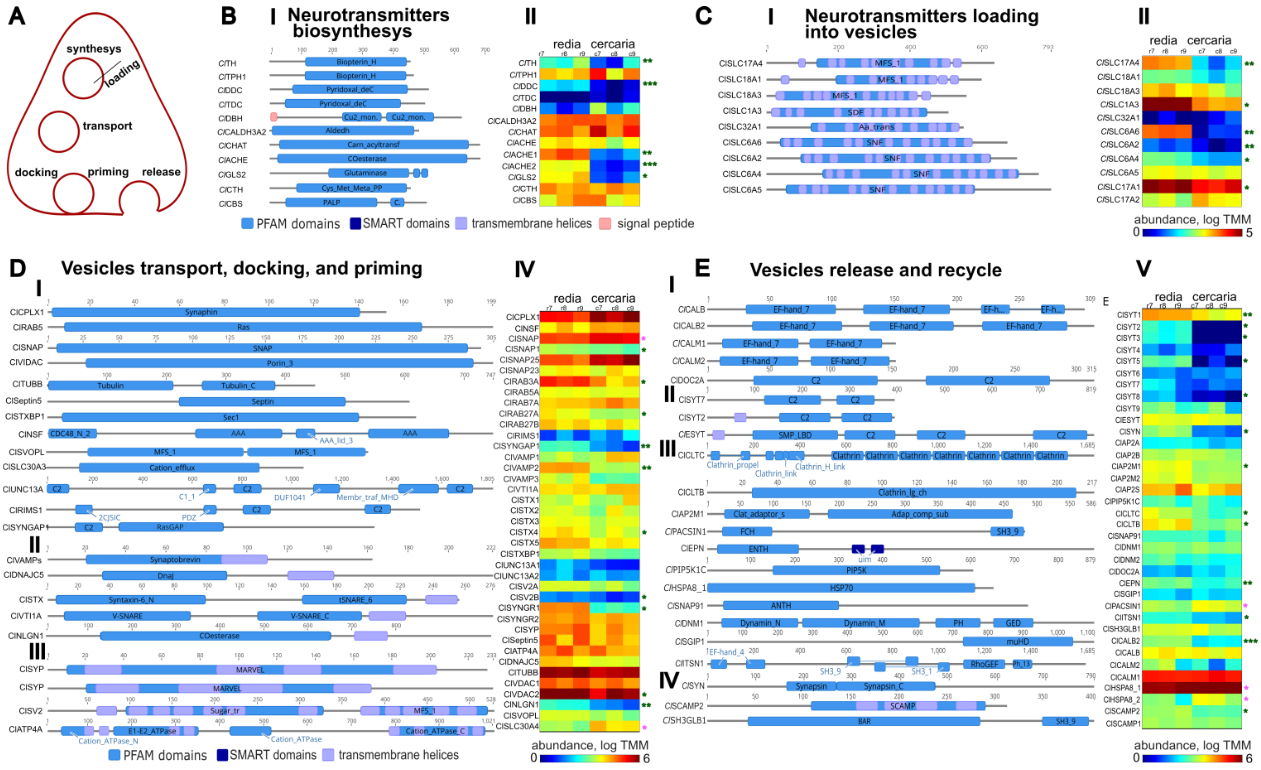


**Supplement figure 1. Vesicle cycle proteins of *C. lingua*.** (**A**) Schematics of vesicle cycle processes. (**B**) Characterization of *C. lingua* proteins related to neurotransmitters biosynthesis and metabolism gene subset. Domain architecture of proteins (**I**) and heat map of the transcript abundance levels (**II**) in rediae and cercariae. Scale in A – protein length (aa). Here and below transcript abundance levels in rediae and cercariae are shown as log of cross-sample normalized TMM; P values: * <0.05; ** <0.01; *** <0.001, green asterix – genes upregulated in redia, statistical test - paired two-tailed T-test. (**C**). Characterization of *C.* lingua proteins related to the subset of packaging and storage of neurotransmitters genes. Domain architecture of the proteins (**I**) and transcript abundance levels in rediae and cercariae (**II**). (**D**) Characterization of *C. lingua* proteins related to transport, docking and priming of synaptic vesicles gene subset. Domain architecture of cytoplasmic proteins (**I**), single-pass membrane proteins (**II**) and multi-pass membrane proteins (**III**). Transcript abundance levels in rediae and cercariae (**IV**). (**E**) Characterization of *C.* lingua proteins related to release and recycle gene subset. Domain architecture of EF-Hand superfamily proteins (**I**), double C2-like domain-containing proteins (**II**), proteins involved in Clathrin-mediated endocytosis (**III**) and proteins associated with small synaptic vesicles (**IV**); scale – protein length (aa). Transcript abundance levels (**V**) in rediae and cercariae. P values: * <0.05; ** <0.01; *** <0.001, green asterix – genes upregulated in redia, magenta asterix – genes upregulated in cercaria, statistical test - paired two-tailed T-test.

*Packaging and storage of neurotransmitters (Synaptic vesicles loading)*

This group of genes encode multi-pass membrane proteins (Supplement Table 2) bearing four characteristic PFAM domains. Major facilitator superfamily (MFS), dicarboxylate symporter family (SDF), sodium:neurotransmitter symporter family (SNF) and amino acid transporter (Aa_trans). glutamate/aspartate transporters in the *C. lingua* transcriptome are represented by five proteins. These are three MSF-proteins, one homolog of vesicular glutamate transporter (*Cl*Slc17A4) and two sialin homologs (*Cl*SLC17A1, 2). SDF-bearing excitatory amino acid transporter (*Cl*SLC1A3) and SNF-bearing sodium- and chloride-dependent taurine transporter (*Cl*SLC6A6) were also classified within the group.

Monoamine transporters are represented by the MSF-bearing protein *Cl*SLC18A1, homologous to Synaptic vesicular amine transporter, and two SNF-bearing proteins: the *Cl*SLC6A2 which is similar to both the sodium-dependent dopamine transporter and the Sodium-dependent norepinephrine transporter, and the *Cl*SLC6A4 homologous to sodium-dependent serotonin transporter. We classified SNF-bearing *Cl*SLC6A5 and Aa_trans-bearing *Cl*SLC32A1 (homolog of Vesicular inhibitory amino acid transporter) as glycine transporters. An MSF-protein *Cl*SLC18A3, the vesicular acetylcholine transporter, was also identified in the *C. lingua* transcriptome (Suppl. Figure 1 C).

*Transport, docking and priming of synaptic vesicles (Vesicle trafficking)*

The first subset, cytoplasmic proteins, includes homologs of complexin (*Cl*CPLX1), alpha-soluble NSF attachment protein (*Cl*SNAP), synaptosomal-associated proteins Munc18 (*Cl*SNAP1), SNAP23, SNAP25 (*Cl*SNAP23,25), septin 5 (*Cl*septin5), syntaxin-binding protein (*Cl*STXBP1), NSF (N-ethylmaleimide sensitive) protein (*Cl*NSF), tubulin-beta (*Cl*TUBB), RAS oncogene family Rab3A, Rab5A, Rab7A, Rab27A,B members (*Cl*RAB3A, *Cl*RAB5A, *Cl*RAB7A, *Cl*RAB27A, *Cl*RAB27B), regulating synaptic membrane exocytosis protein 1 (*Cl*RIMS1), Ras/Rap GTPase-activating protein SynGAP (*Cl*SYNGAP1), two homologous genes of voltage-dependent anion-selective channel (*Cl*VDAC1, 2), two Munc13a homologs (*Cl*UNC13A1, 2), putative transporter SVOPL (*Cl*SVOPL) and synaptic vesicle associated Zinc transporter (*Cl*SLC30A4). The second, single-pass membrane proteins, consists of three homologs of vesicle-associated membrane protein (*Cl*VAMP1-3), homolog of cysteine string protein (*Cl*DNAJC5), vesicle transport through interaction with t-SNAREs homolog (*Cl*VTI1A), five genes encoding syntaxins (*Cl*STX1-5) and neuroligin-1 homolog (*Cl*NLGN1). The third subset, multi-pass membrane proteins, includes synaptophysin (*Cl*SYP), two synaptogyrins (*Cl*SYNGR1, 2), sodium/potassium-transporting ATPase subunit alpha (*Cl*ATP4A) and two chains of synaptic vesicle glycoprotein 2 (*Cl*SV2A, B) (Supplement table 3, Suppl. Figure 1 D).

*Vesicles release and recycle*

We classified four EF-Hand superfamily proteins found in the transcriptome of *C. lingua* as calbindin (*Cl*CALB), calretinin/Calbindin2 (*Cl*CALB2) and two variants of calmodulin (*Cl*CALM1, 2). The family of double C2-like domain-containing proteins is represented by the ortholog of DOC2A (*Cl*DOC2A) and 10 homologs of synaptotagmin (*Cl*SYT1-9, *Cl*ESYT). Among synaptotagmins, four proteins are cytoplasmic (*Cl*SYT6-9) and other six are membrane anchored including one homolog of extended synaptotagmin (*Cl*ESYT). Proteins of clathrin-associated complex involved in clathrin-mediated endocytosis are rather diverse in domain architecture. They are represented with clathrin heavy (*Cl*CLTC) and light (*Cl*CLTB) chains, clathrin coat assembly protein (*Cl*SNAP91), epsin (*Cl*EPN), two dynamin homologs (*Cl*DNM1,2), intersectin-1 (*Cl*ITSN1), phosphatidylinositol 4-phosphate 5-kinase type-1 gamma (*Cl*PIP5K1C), SH3-containing GRB2-like protein 3-interacting protein 1 (*Cl*SGIP1), AP-2 complex subunits (*Cl*AP2A1, B1, S1,M1,M2), two homologs of co-chaperone Heat shock cognate 71 kDa protein (ClHSPA8_1, 2) and protein kinase C/casein kinase substrate neurons (*Cl*PACSIN1). In Platyhelminthes the homolog of the gene PACSIN1 is annotated as Antigen EG13. Finally, proteins associated with small synaptic vesicles, synapsin (*Cl*SYN), two homologs of secretory carrier-associated membrane protein 1 (*Cl*SCAMP1, 2) and SH3-domain GRB2-like endophilin B1 (*Cl*SH3GLB1) (Supplement table 4, Suppl. Figure 1 E).

*Reception machinery*

Neurotransmitter ligand-gated ion channels are transmembrane receptor-ion channel complexes and contain receptor family ligand binding region and neurotransmitter-gated ion-channel transmembrane region. These channels open transiently upon binding of specific ligands, allowing rapid transmission of signals at chemical synapses. We classified nicotinic acetylcholine receptors *Cl*ACC-1.1, *Cl*ACC-1.2, *Cl*ACC-2, *Cl*ACC-2.1, *Cl*ACC-2.2, *Cl*CHRACN1, *Cl*CHRACN2, *Cl*CHRACN3, *Cl*CHRACN4 and ClCHRACN5 orthologs of the Acetylcholine gated chloride channels of *Schistosoma mansoni* ACC-1 and ACC-2 in the transcriptome studied. We also identified another inhibitory chloride ion channel glycine receptors, both alpha and beta subunits *Cl*GLRA, *Cl*GLRA2, *Cl*GLRB. From this group of receptors we also classified glutamate receptors, containing ligated ion channel L-glutamate-binding site excitatory cation channels kainite and N-methyl-D-aspartate (NMDA) - *Cl*GRIK1, *Cl*GRIK2, *Cl*GRIK3, *Cl*GRIK4, *Cl*GRID2, *Cl*GRIN1 and chloride channel *Cl*GLUCL.

Metabotropic receptors contain ligand binding ANF-receptor and seven transmembrane domains. We identified several putative g-protein coupled receptor families including orthologs of muscarinic acetylcholine receptors *Cl*GAR1 and *Cl*GAR2, orthologs of dopamine receptors *Cl*D2R1, *Cl*D2R2, orthologs of serotonin receptors *Cl*HTR1A1-A5, octopamine-tyramine receptors *Cl*Octbeta1 and *Cl*Oct2, and ortholog of metabotropic glutamate receptor (GRM2-5) *Cl*GRM1. We also identified ortholog of Inositol 1,4,5-trisphosphate receptor type 1 *Cl*ITPR1,2 containing ion transport domain.

A portion of reception machinery in *C. lingua* transcriptome is represented by cytoplasmic proteins. The *Cl*SHANK1 contain Ferm Fo, ankyrin and PDZ domains and represents an ortholog of an adapter protein SHANK1 that interconnects metabotropic glutamate receptors and the actin-based cytoskeleton. The *Cl*DLG1 is an ortholog of the Disks large homolog (DLG1-4) playing role in synaptogenesis and chemical synaptic transmission and contain a receptor targeting L 27, PDZ and guanylate kinase domains. The *Cl*HOMER1 is a ortholog of homer protein (HOMER1,3) -- postsynaptic density scaffolding protein regulating synaptic metabotropic glutamate function and contain WASP homology region 1. *Cl*DBNL is an ortholog of drebrin-like protein (DBN1) playing a role in neuron morphogenesis and synapse formation. It contains actin depolymerisation factor/cofilin -like and Src homology 3 domains. The *Cl*STRN1 is an ortholog of striatin (STRN), a calmodulin-binding protein which may function as scaffolding or signaling protein and may play a role in dendritic Ca2+ signaling and contain eponymous domain. We also identified the *Cl*DLGAP1 containing an eponymous domain an ortholog of disks large-associated protein (DLGAP1-4) which is a part of the postsynaptic scaffold in neuronal cells. We also classified orthologs of P2X purinoreceptor ClP2X1, ClP2X2 containing eponymous domain and of guanylate cyclase soluble subunit beta-1 containing NO binding HNOB domain (Supplement table 5, Suppl. Figure 2A).

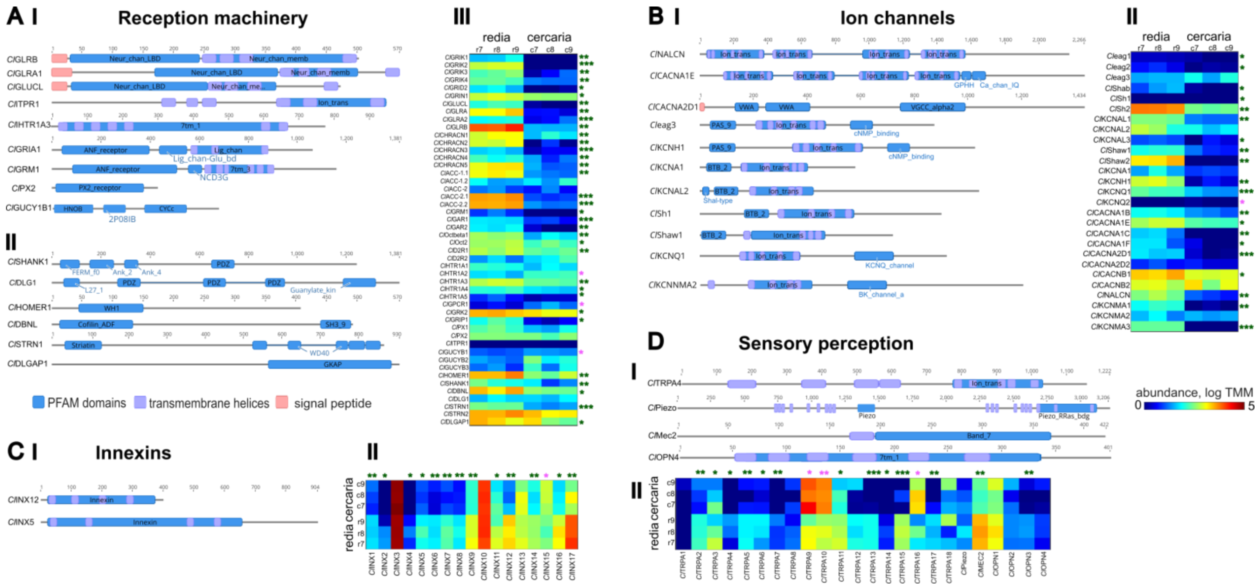


**Supplement figure 2**. (**A**) **Characterization of *C.* lingua proteins related to reception machinery gene subset.** Domain architecture of representative multi-pass membrane proteins (**I**) and cytoplasmic proteins (**II**); scale – protein length (aa). Transcript abundance levels in rediae and cercariae (**III**). (**B**). Characterization of *C. lingua* ion channels gen subset. Domain architecture of sodium leak channel, potassium and calcium voltage gated channels (**I**) and their expression abundance in rediae and cercariae (**II**). (**C**) Characterization of *C. lingua* innexins. Domain architecture of innexins (**I**) and their expression profiles (**II**). (**D**) Characterization of *C. lingua* sensory gene subset. Domain architecture of proteins (**I**) and their transcript abundance in rediae and cercariae (**II**). Scale – protein length (aa). P values: * <0.05; ** <0.01; *** <0.001, green asterix – genes upregulated in redia, magenta asterix – genes upregulated in cercaria, statistical test - paired two-tailed T-test.

*Ion channels*

Of all $Na^+$ channel proteins only a sodium leak channel non-selective protein (*Cl*NALCN) was identified. Domain architectures of *C. lingua* voltage-dependent calcium channels, R-type subunit alpha-1E (*Cl*CACNA1E) N-type subunit alpha-1B (*Cl*CACNA1B), L-type subunit alpha-1C (*Cl*CACNA1C) and L-type subunit alpha-1F (*Cl*CACNA1F) are similar, hallmarked with C-terminal domains GPHH, Ca_chan_IQ and shown in Figure 12 C (*Cl*CACNA1E). The alpha-2/delta subunits of voltage-dependent calcium channels (*Cl*CACNA2D1 and *Cl*CACNA2D2) are characterized with two N-terminal von Willebrand factor (vWF) type A domains and a specific C-terminal domain VGCC_alpha2.

We identified some of *C. lingua* potassium voltage-gated channels , encoding three orthologs of protein eag (Cleag1 - 3), subfamily H member (*Cl*KCNH1), subfamily A member (*Cl*KCNA1), three proteins Shal (*Cl*KCNAL1 - 3) with characteristic N-terminal domain Shal-type, two proteins Shaker (*Cl*Sh1, 2) homologues to functionally characterized SKv1.1 channel, protein Shab (*Cl*Shab), two proteins Shaw (*Cl*Shaw1, 2), two members of KQT-subfamily (*Cl*KCNQ1, 2), four transmembrane inward-rectifying TWIK-related potassium channel (*Cl*KCNK9) and three potassium/sodium hyperpolarization-activated cyclic nucleotide-gated channels (*Cl*HCN1-3). Among three types of Ca2+-activated K+ channel we classified homologs of small-conductance (SK) type characterized by the SK channel and Ca2+-binding protein calmodulin (CaM) domain (*Cl*KCNN1-3) and large conductance type with BK channel a domain (*Cl*KCNMA1-3). We also classified homologs of cGMP-gated cation channel subunits (*Cl*CNGA1-3, *Cl*CNGB) (Supplement table 6, Suppl. Figure 2B).

*Innexins*

Innexins forming gap junctions and non-junctional membrane channels were found expressed in *C. lingua.* They bear a hallmark domain with four transmembrane helices. Innexins were represented in the transcriptome by 16 genes (*Cl*INX1 - 16). Only two of them were expressed at levels above 100 TMMs (Supplement table 7, Suppl. Figure 2C).

*Sensory perception proteins*

We identified orthologs of transient receptor ion channels *Cl*TRPA1-A18 containing transient receptor ion channel, polycystin cation channel, ion transport protein domains. We classified ortholog of another non-specific cation channel, Piezo mechanosensitive ion channel (*Cl*Piezo) containing eponymous domains and ortholog of mechanosensory protein 2 mec-2 containing prohibitin homologues domain *Cl*MEC2 . We also identified orthologs of light-absorbing opsins *Cl*OPN1-4 from the ion transport protein members of the seven-transmembrane-domain proteins of the G protein-coupled receptor (GPCR) superfamily (Supplement table 8, Suppl. Figure 2D).
